# Amyloid-β fibrils accumulated in preeclamptic placentas suppress syncytialization of cytotrophoblasts

**DOI:** 10.1101/2025.03.06.641957

**Authors:** Kaho Nishioka, Midori Ikezaki, Naoyuki Iwahashi, Miyu Arakawa, Momo Fukushima, Noa Mori, Mika Mizoguchi, Yuko Horiuchi-Tanizaki, Megumi Fujino, Takami Tomiyama, Yoshito Ihara, Kenji Uchimura, Kazuhiko Ino, Kazuchika Nishitsuji

**Author notes:** Corresponding author: Kazuchika Nishitsuji, Ph.D., Department of Biochemistry, Wakayama Medical University School of Medicine, 811-1 Kimiidera, Wakayama 641-8509, Japan.

## Abstract

Cerebral deposition of fibrillar amyloid-β (Aβ) is a pathological hallmark of Alzheimer’s disease. While Aβ is present in human placentas and accumulates in preeclamptic placentas characterized by poor placentation, the production and role of Aβ in the human placenta remain unclear. Because hypoxia in mid-to-late pregnancy is a risk for preeclampsia, we found that levels of hypoxia-inducible factor 1-α and β-secretase (BACE-1) increased concurrently with placental Aβ deposition in late stage preeclamptic placentas. We also found that a human cytotrophoblast (CTB) model, BeWo cells, actually produced Aβ species, and that hypoxia increased Aβ production and BACE-1 protein levels. Aβ42 fibrils inhibited CTB syncytialization, a critical step in maintaining pregnancy, by inducing loss of membrane localization of cell-cell adhesion molecules. Primary human CTBs confirmed these observations. Taken together, our results suggest that increased Aβ production in CTBs by hypoxia may lead to the formation of Aβ fibrils, which inhibit syncytiotrophoblast formation and are detrimental to pregnancy. Thus, our results reveal the novel role of Aβ fibrils in the pathogenesis of preeclampsia.

## INTRODUCTION

Preeclampsia (PE) is one of the most severe pregnancy-specific disorders associated with hypertension and proteinuria. Approximately 7–10% of pregnancies manifest PE, which may result in high maternal and fetal morbidity and mortality (Abalos, Cuesta, Grosso, Chou, & Say, 2013; Duley, 2009; Khan, Wojdyla, Say, Gulmezoglu, & Van Look, 2006; Wallis, Saftlas, Hsia, & Atrash, 2008). At this time, no cure other than delivery of the fetus exists for PE, which makes PE the leading cause of iatrogenic preterm birth (Phipps, Thadhani, Benzing, & Karumanchi, 2019). Although the etiology of PE is not yet fully understood, poor placentation has been implicated in PE pathophysiology (Aplin, Myers, Timms, & Westwood, 2020; Chappell, Cluver, Kingdom, & Tong, 2021). Several studies reported deposition of misfolded proteins in PE placentas (Buhimschi et al., 2014; Cater et al., 2019). These proteins include amyloid-β (Aβ) peptides, which deposit in the brains of patients with Alzheimer’s disease (AD). Aβ is produced by the sequential cleavage of amyloid precursor protein (APP) by β-secretase 1 (BACE1) and γ-secretase. Aβ levels in the brain are determined by the balance between Aβ production and clearance (Selkoe & Hardy, 2016). Thus, an imbalance between Aβ production and clearance leads to increased Aβ levels as well as Aβ aggregation to form toxic Aβ aggregates. APP is widely expressed throughout the body, including the placenta, and can be processed by BACE1 and γ-secretase to produce Aβ in the placenta (Buhimschi et al., 2014; Mattson, 2004). Although Aβ aggregates have been established as toxic to neurons (Selkoe & Hardy, 2016), exactly how Aβ and Aβ aggregates affect placental cell functions is unknown.

Cytotrophoblasts (CTBs) are epithelial stem cells in the human placenta that differentiate into two major placental cell types: extravillous trophoblasts (EVTs) and syncytiotrophoblasts (STBs) (Bischof & Irminger-Finger, 2005). CTBs undergo continuous syncytialization to form STBs in the outer layers of the floating chorionic villi, which are critical for key placental functions such as fetal nutrition, gas exchange, and protection and placental hormone production (Eaton & Contractor, 1993; Kliman, Nestler, Sermasi, Sanger, & Strauss, 1986; Ogren & Talamentes, 1994). Hypoxia reportedly inhibited CTB syncytialization via a transcription factor complex known as hypoxia-inducible factor (HIF) (Jaremek et al., 2023), and CTB syncytialization was reduced in PE placentas (Costa, 2016), which indicated a role of hypoxia in poor placentation and PE pathology.

Hypoxia associated with cerebral ischemia and stroke is a strong risk for the development of late-onset AD (Kokmen, Whisnant, O’Fallon, Chu, & Beard, 1996; Tao et al., 2024; Tatemichi et al., 1994). Hypoxia associated with cerebral ischemia and stroke is thought to increase Aβ production by inducing expression of BACE1 (Sun et al., 2006; X. Zhang et al., 2007). Because PE and AD share hypoxia as a common factor that is implicated in the pathogenesis, we hypothesized that hypoxia may increase Aβ production in preeclamptic placentas, leading to the formation of toxic fibrillar Aβ and inhibiting STB formation. The main Aβ peptide species are Aβ40 and Aβ42, and Aβ42 is more predisposed to aggregation than is Aβ40 (Tanzi & Bertram, 2005). Indeed, genetically modified mice that generate Aβ42, but not Aβ40 alone, developed amyloid plaques (McGowan et al., 2005). Therefore, chronic hypoxia in the dysplastic placenta may increase Aβ production and subsequent formation of toxic Aβ aggregates. Here, we report Aβ deposition in preeclamptic placentas and that hypoxia enhanced BACE1 expression and Aβ production in CTB model BeWo cells. We also show that Aβ fibrils suppressed differentiation of CTB model BeWo cells and primary human CTBs. Our results support the pathological role of Aβ fibrils in poor placentation by interfering with STB formation.

## RESULTS

### Aβ accumulated and hypoxia was enhanced in PE placentas

We performed immunohistochemical analyses of normal and PE placentas in late pregnancy with the β001 anti-amyloid β antibody. The information for each of the cases is shown in Table 1. Of the 5 PE cases, one was a late-onset case and one was an early-onset case diagnosed with intrauterine growth restriction. We used the RB4CD12 anti-heparan sulfate S-domain antibody as an amyloid/protein aggregate marker, because heparan sulfate S-domains have been shown to co-deposit with amyloid *in vivo* (Bruinsma et al., 2010; Hosono-Fukao et al., 2012; Iwahashi et al., 2020; Kameyama et al., 2019; Kazuchika Nishitsuji, 2018; K. Nishitsuji & Uchimura, 2017). In the villi of 5 PE cases, but not in normal placentas, we found co-deposition of Aβ with the RB4CD12 epitope (Fig. 1). The transcription factor HIF-1α is activated in response to hypoxia and plays an important role in cell responses and adaptation to hypoxia (Weidemann & Johnson, 2008). Hypoxia and HIF1-α reportedly enhanced BACE1 expression and Aβ production (Sun et al., 2006; X. Zhang et al., 2007). Because PE placentas exist under hypoxic conditions (Tong et al., 2022), we analyzed the expression of HIF1-α and BACE1 in human normal and PE placentas. Immunohistochemistry revealed induction of HIF-1α in PE placentas and BACE1 expression around HIF-1α-positive cells in chorionic villi (Fig. 2). HIF-1α was mainly induced in STBs and CTBs, whereas BACE1 expression was observed in CTBs and STBs as well as in stromal tissues in PE placental villi. Signal intensities of HIF1-α in normal placentas were weak and significantly lower than those of PE placentas, which was consistent with a previous report (Caniggia & Winter, 2002).

**Fig. 1.**
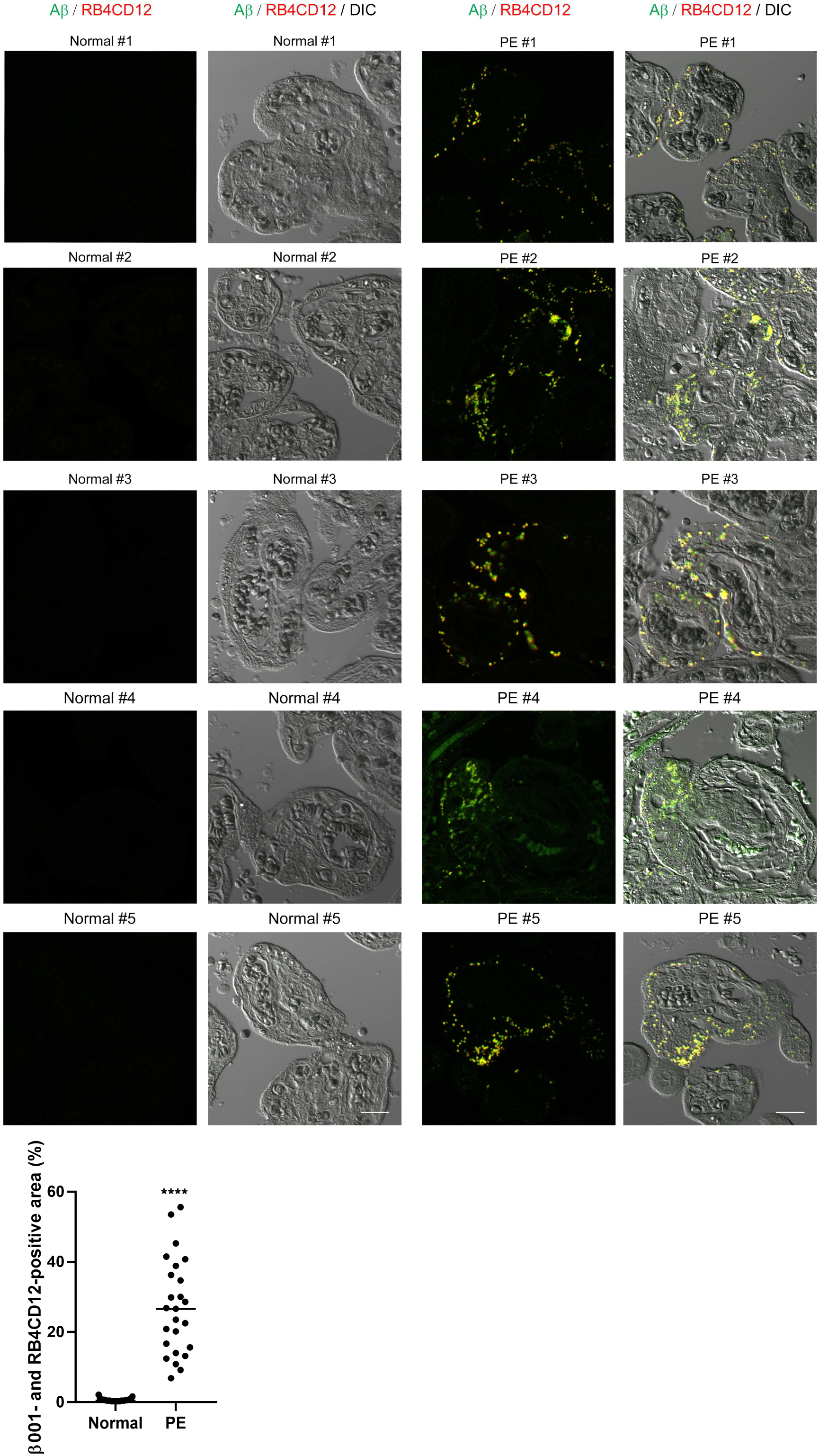
Aβ deposition in PE placentas. Immunohistochemical analysis of normal and PE human placentas from the third trimester. Sections were immunostained with the β001 rabbit polyclonal anti-amyloid β antibody and RB4CD12, a marker of co-deposition of amyloid/protein aggregates. Graph shows the quantification of β001- and RB4CD12-positive area in ROIs. \*\*\*\**P* < 0.0001. Scale bar, 20 µm.

**Figure 2.**
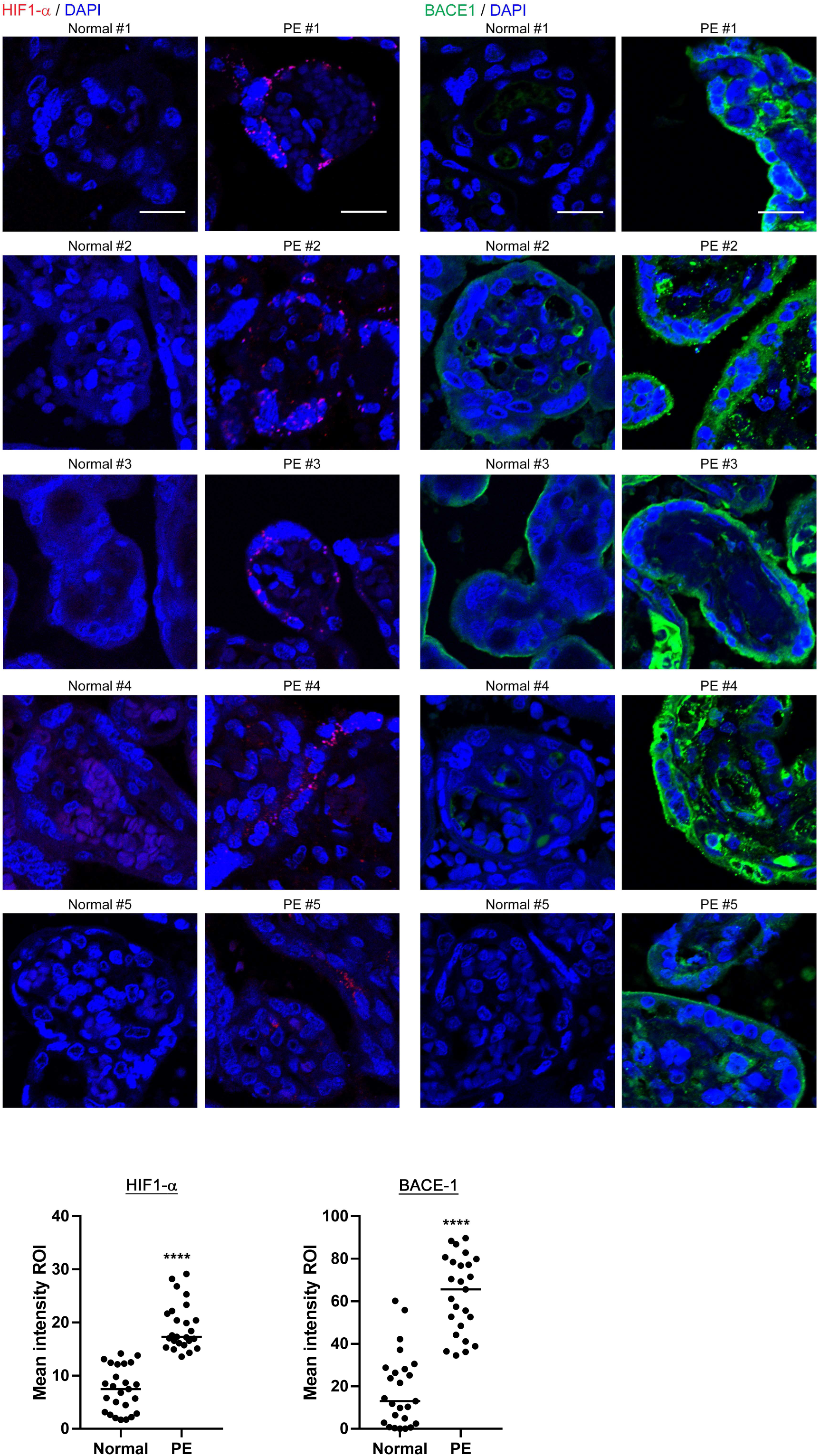
HIF1-α and BACE1 levels in PE placentas. Sections were immunostained with the anti-HIF-1α antibody or the anti-BACE1 antibody. Nuclei were counterstained with DAPI. Graphs show the quantification of the mean intensities of HIF1-α and BACE-1 signals. \*\*\*\**P* < 0.0001. Scale bars, 20 µm.

**Table 1.**
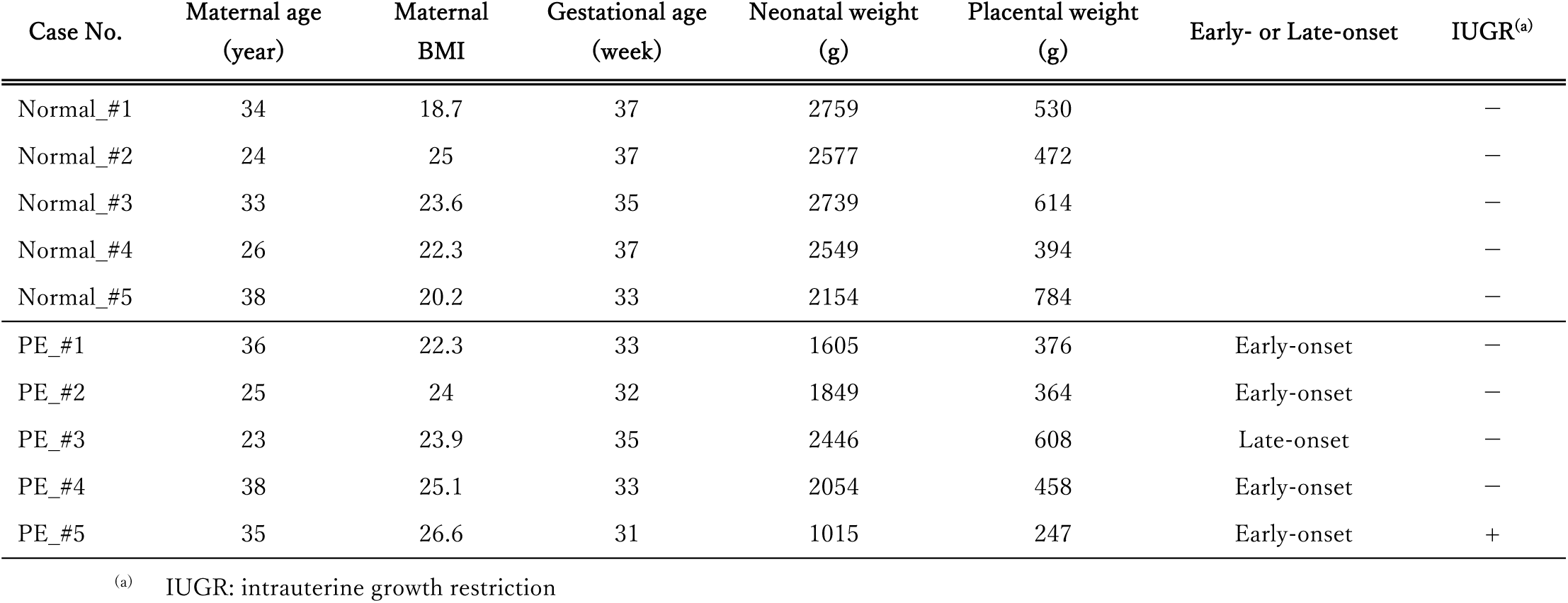
Clinical information of the study population.

| Case No. | Maternal age<br>(year) | Maternal<br>BMI | Gestational age<br>(week) | Neonatal weight<br>(g) | Placental weight<br>(g) | Early- or Late-onset | IUGR<sup>(a)</sup> |
| --- | --- | --- | --- | --- | --- | --- | --- |
| Normal_#1 | 34 | 18.7 | 37 | 2759 | 530 |  | — |
| Normal_#2 | 24 | 25 | 37 | 2577 | 472 |  | — |
| Normal_#3 | 33 | 23.6 | 35 | 2739 | 614 |  | — |
| Normal_#4 | 26 | 22.3 | 37 | 2549 | 394 |  | — |
| Normal_#5 | 38 | 20.2 | 33 | 2154 | 784 |  | — |
| PE_#1 | 36 | 22.3 | 33 | 1605 | 376 | Early-onset | — |
| PE_#2 | 25 | 24 | 32 | 1849 | 364 | Early-onset | — |
| PE_#3 | 23 | 23.9 | 35 | 2446 | 608 | Late-onset | — |
| PE_#4 | 38 | 25.1 | 33 | 2054 | 458 | Early-onset | — |
| PE_#5 | 35 | 26.6 | 31 | 1015 | 247 | Early-onset | + |

<sup>(a)</sup> IUGR: intrauterine growth restriction

### Hypoxia increased BACE1 levels and Aβ production in CTB model cells

In our study, we used BeWo cells, which are widely used as a model for trophoblast syncytialization (Drewlo, Baczyk, Dunk, & Kingdom, 2008; Gauster, Moser, Orendi, & Huppertz, 2009). Prolyl hydroxylation of HIF1-α acts as a signal for the ubiquitin-proteasome mediated degradation of HIF1 (Schofield & Ratcliffe, 2004). In order to study the effect of HIF-1α on BACE1 expression, we used Roxadustat, which inhibits the prolyl hydroxylation and subsequent degradation of HIF1-α (Hsieh et al., 2007). Roxadustat (0, 5, and 10 μM) significantly increased HIF-1α expression (3.1-fold for 5 μM versus 0 μM, and 5.5-fold and 1.5-fold for 10 μM versus 0 μM and 5 μM, respectively) and BACE1 (1.4-fold for 5 μM and 1.6-fold for 10 μM versus 0 μM) in BeWo cells (Fig. 3A). Thus, hypoxia-induced up-regulation of HIF-1α enhanced BACE1 expression in BeWo cells, which may eventually increase Aβ production.

**Figure 3.**
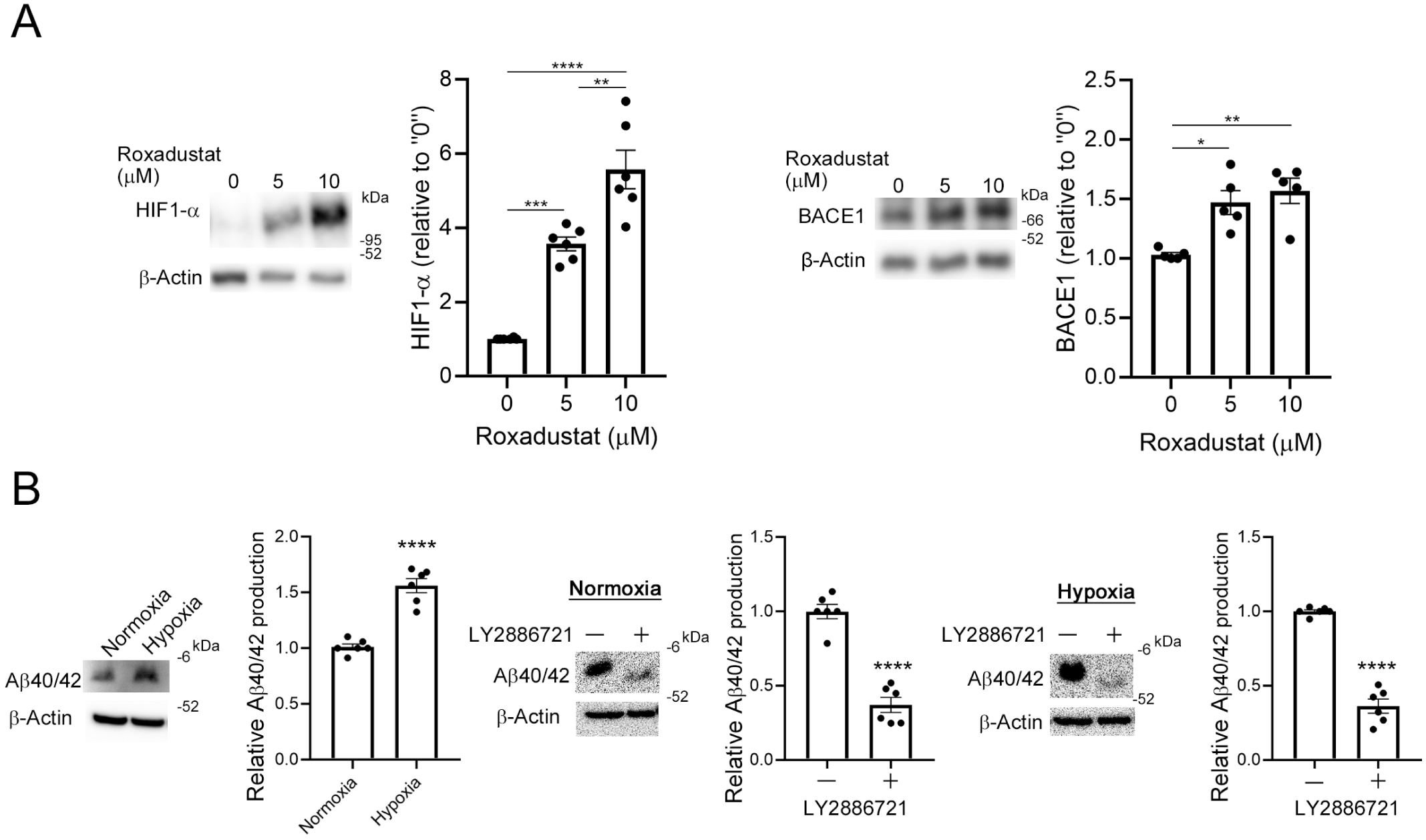
(**A**) BeWo cells were cultured in 15% MEM-α with roxadustat (0, 5, or 10 μM) for 16 hours. Protein levels of HIF-1α and BACE1 were analyzed by means of Western blotting with the anti-HIF-1α antibody and the anti-human BACE1 antibody, respectively. β-Actin served as the loading control. Graphs show quantification of HIF-1α and BACE1. Data are means ± SEM (*n* = 6). \**P* < 0.05; \*\**P* < 0.01; \*\*\**P* < 0.001; \*\*\*\**P* < 0.0001. (**B**) Hypoxia promoted Aβ production in BeWo cells. Aβ40/42 generated by BeWo cells was analyzed by means of Western blotting with the anti-Aβ (Nt) β001 antibody. BeWo cells were cultured in Opti-MEM medium containing 2% FBS with or without LY2886721 for 24 hours under normoxic conditions (20% O_2_) or hypoxic conditions (2% O_2_). Generated Aβ40/42 was analyzed. β-Actin served as the loading control. Graphs show quantification of Aβ40/42. Data are means ± SEM (*n* = 6). \*\*\**P* < 0.001; \*\*\*\**P* < 0.0001.

We then analyzed Aβ production in these cells by means of Western blotting with the β001 anti-Aβ N-terminus (Nt) antibody (Lippa et al., 1999). Aβ40/42 production in BeWo cells increased 150% under hypoxic conditions compared with that under normoxic conditions (Fig. 3B). We also found that the LY2886721 BACE1 inhibitor reduced Aβ40/42 production in BeWo cells under normoxic and hypoxic conditions (40% and 33%, respectively; Fig. 3B). We did not observe Aβ oligomers in CM obtained from BeWo cells (Supplementary Figure S1).

### Aβ42 fibrils inhibited syncytialization of CTB model BeWo cells

Hypoxia has been implicated in placentation dysfunction after the formation of spiral arteries. We therefore hypothesized that increased production of Aβ by CTBs and subsequent aggregation of Aβ may have detrimental effects on CTB functions and thereby contribute to defects in placentation and development. Although Aβ40 is generally the predominant specie (Burdick et al., 1992), Aβ42 is more prone to aggregation than Aβ40 (Suh & Checler, 2002). Because syncytialization of CTBs is a critical event for placentation, is maintained until the end of pregnancy, and is reportedly impaired in PE placentas (Costa, 2016), we investigated the effect of fibrillar Aβ42 on forskolin (Fsk)-induced syncytialization by analyzing the secretion and induction of human chorionic gonadotropin β-subunit (β-hCG) (Gerbaud & Pidoux, 2015; Wice, Menton, Geuze, & Schwartz, 1990) and induction of syncytin-1 expression (Mi et al., 2000) in BeWo cells. Here, pre-treatment of BeWo cells with Aβ42 fibrils in a micromolar range significantly reduced the secretion and induction of β-hCG by 40% in the medium and reduced the expression of syncytin-1 by 65% (Fig. 4A).

**Figure 4.**
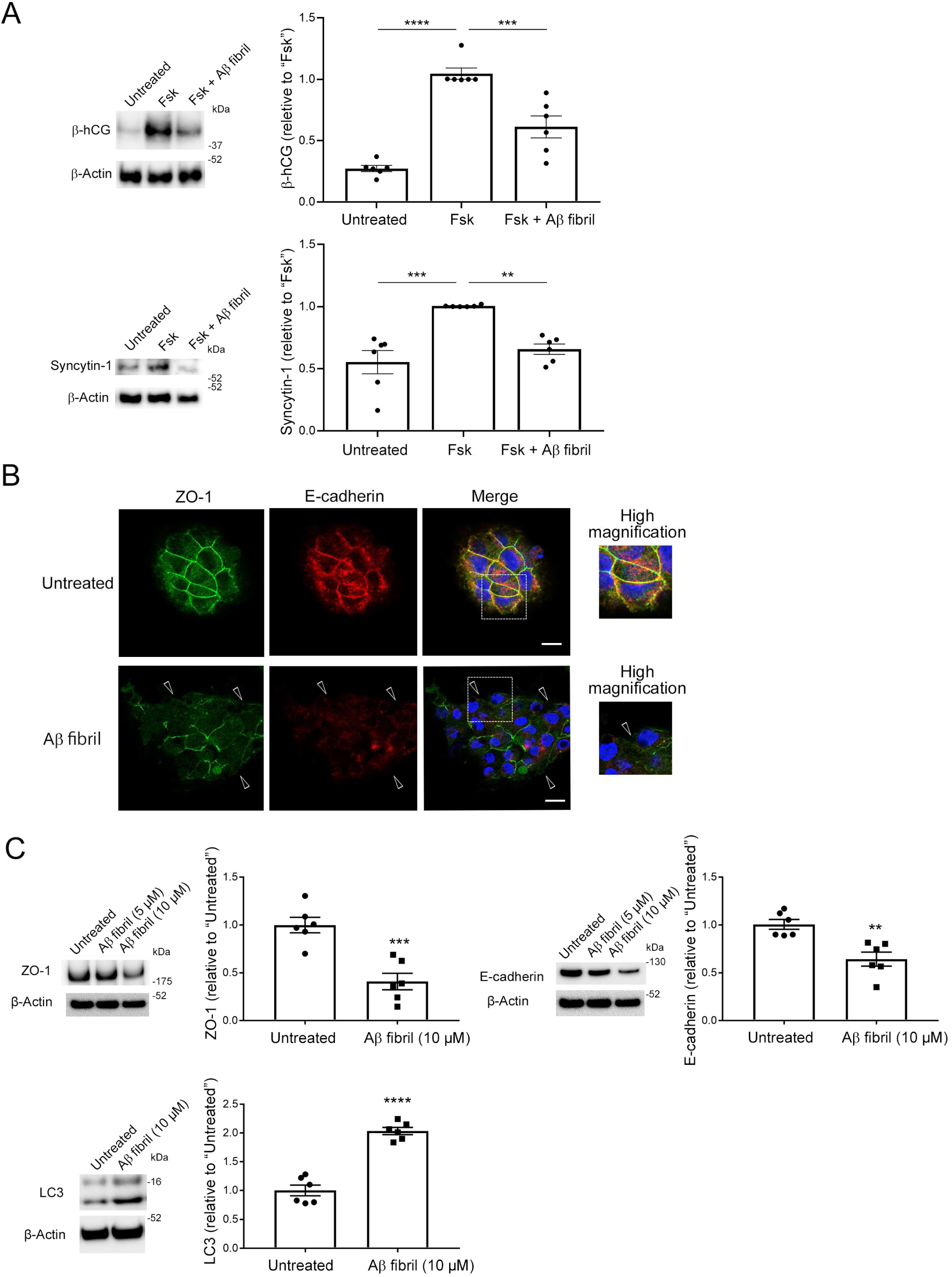
Aβ42 fibrils inhibited syncytialization of BeWo cells by inducing loss of membrane localization of cell-cell adhesion proteins. (**A**) BeWo cells were pre-treated with Aβ42 fibrils (10 μM) in serum-free Opti-MEM for 12 hours and then were stimulated with Fsk (50 μM) for 48 hours. The effect of Aβ fibrils on syncytialization of BeWo cells was analyzed by means of Western blotting with the anti-hCG β antibody and the anti-syncytin-1 antibody. β-Actin served as the loading control. Graphs show quantification of β-hCG and syncytin-1. Data are means ± SEM (*n* = 6). \*\**P* < 0.01; \*\*\**P* < 0.001; \*\*\*\**P* < 0.0001. (**B**) BeWo cells were cultured on cover glasses and treated with Aβ42 fibrils (10 μM) for 24 hours, after which they were fixed in 4% paraformaldehyde and stained with the anti-ZO-1 antibody or the anti-E-cadherin antibody. Arrowheads indicate loss of membrane localization of ZO-1 and E-cadherin in Aβ fibril-treated BeWo cells. Nuclei were counterstained with DAPI. Scale bars, 20 µm. (**C**) BeWo cells were treated with Aβ42 fibrils (10 μM) in serum-free Opti-MEM for 24 hours. Quantitative analysis of ZO-1 and E-cadherin of Aβ42 fibril-treated BeWo cells was performed by using Western blotting with the anti-ZO-1 antibody and the anti-E-cadherin antibody. Evaluation of the effect of Aβ42 fibrils on autophagy of BeWo cells was performed by using Western blotting with an anti-LC3 antibody. β-Actin served as the loading control. Graphs show quantification of ZO-1, E-cadherin, and LC3. Data are means ± SEM (*n* = 6). \*\**P* < 0.01; \*\*\**P* < 0.001; \*\*\*\**P* < 0.0001.

Cell-cell adhesion proteins such as ZO-1 and E-cadherin are required for CTB syncytialization (Iwahashi et al., 2019; Pidoux et al., 2010). Therefore, we next investigated the effect of Aβ42 fibrils on subcellular localization of ZO-1 and E-cadherin in BeWo cells. Immunofluorescence analysis with an anti-ZO-1 antibody and an anti-E-cadherin antibody revealed that Aβ42 fibrils disrupted the membrane localization of ZO-1 and E-cadherin (Fig. 4B, arrowheads). Immunoblots showed significant reductions in ZO-1 and E-cadherin protein levels in Aβ42 fibril-treated BeWo cells (34% for ZO-1 and 45% for E-cadherin, Fig. 4C). A previous study suggested that Aβ fibril treatment enhanced autophagy and thereby reduced levels of cell adhesion-related proteins including ZO-1 in endothelial cells (Chan et al., 2018). Turnover of E-cadherin is at least partly regulated via the autophagic pathway (Santarosa & Maestro, 2021). Here, Aβ42 fibril-treated BeWo cells showed a significant 260% increase in LC3 levels, which suggested that increased autophagy resulted in decreases in and loss of membrane localization of ZO-1 and E-cadherin (Fig. 4C). Although Aβ42 fibrils increased the mRNA expression of E-cadherin (Supplementary Figure S2), these results suggest that Aβ42 fibrils disrupted the proper membrane localization of cell adhesion-related proteins and thereby interfered with Fsk-induced syncytialization. We also excluded the possibility that Aβ fibrils induced cell death in BeWo cells (Supplementary Figure S3).

### Aβ fibrils inhibited syncytialization of human primary cultured CTBs

Because BeWo cells require Fsk for syncytialization and human CTBs spontaneously undergo syncytialization without Fsk (Costa, 2016), we further investigated the effect of Aβ fibrils on human placentation by using primary cultured human trophoblast cells. We isolated trophoblast cell fractions from human normal placentas as previously described (Simon, Bucher, & Maloyan, 2017). These cells secreted β-hCG 72 hours after seeding, which suggested that CTBs were successfully isolated (Fig. 5A). We also confirmed by means of Western blotting that human primary cultured CTBs produced Aβ40/42. We pre-treated human CTBs with Aβ42 fibrils (10 µM) for 24 hours, after which the culture media were replaced with fresh media containing Aβ42 fibrils (10 µM), and incubation continued for an additional 48 hours. We then analyzed the induction and secretion of β-hCG and induction of syncytin-1 by means of Western blotting, which revealed that treatment of human CTBs with Aβ42 fibrils significantly reduced β-hCG induction and secretion—38% and 65%, respectively (Fig. 5B). Aβ42 fibrils also reduced syncytin-1 induction by 65%, which was similar to the findings for BeWo cells (Fig. 5B). We then asked whether Aβ42 fibrils would affect subcellular localization of ZO-1 and E-cadherin in human CTBs. Immunocytochemical analysis of Aβ42 fibril-treated human trophoblasts revealed disrupted membrane localization of ZO-1 and E-cadherin (Fig. 5C, arrowheads). These results strongly support the finding that Aβ fibrils disrupted the proper localization of ZO-1 and E-cadherin at the cell-cell border and thereby inhibited syncytialization of CTBs.

**Figure 5.**
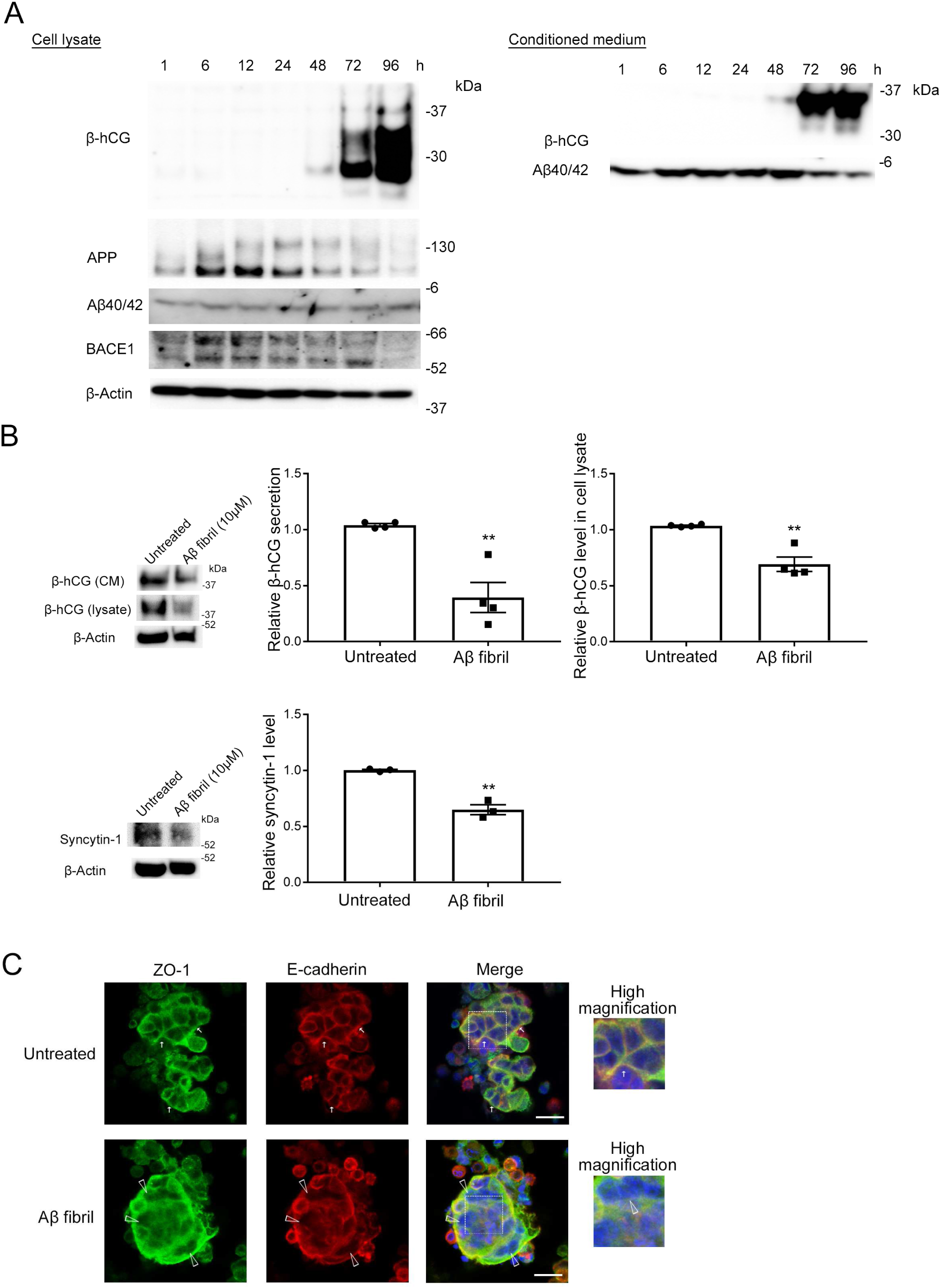
Aβ42 fibrils inhibited primary cultured human CTB syncytialization. (**A**) Primary cultured human CTBs ware isolated from third-trimester human placentas and cultured in Trophoblast Medium supplemented with 5% FBS for 96 hours, after which the expression of APP and BACE1 and the production and secretion of β-hCG and Aβ40/42 were analyzed by means of Western blotting. (**B**) Primary cultured human CTBs were treated with Aβ1-42 fibrils (10 μM) in serum-free Opti-MEM for 72 hours. The effect of Aβ fibrils on syncytialization of human CTBs was analyzed by means of Western blotting with the anti-hCG β antibody and the anti-syncytin-1 antibody. β-Actin served as the loading control. Graphs show quantification of β-hCG and syncytin-1. Data are means ± SEM (*n* = 3 or 4). \*\**P* < 0.01. (**C**) Primary cultured human CTBs were cultured on cover glasses and were treated with Aβ1-42 fibrils (10 μM) for 18 hours, after which they were fixed in 4% paraformaldehyde. They were then stained with the anti-ZO-1 antibody or the anti-E-cadherin antibody. Arrows indicate ZO-1 and E-cadherin located at the cell-cell border, and arrowheads indicate loss of membrane localization of these proteins. Nuclei were counterstained with DAPI. Scale bars, 20 µm.

## DISCUSSION

PE is a complicated syndrome with multifactorial pathology, whose etiology is not well understood. Given that the current definitive treatment of PE is termination of pregnancy, elucidating the mechanisms of placentation and placental defects is important for prevention of PE and development of novel PE therapies. Numerous studies have suggested that the aberrant accumulation of misfolded proteins and their aggregates underlies the pathology of human diseases such as AD, Parkinson’s disease, age-related macular degeneration, arthritis, and p53-mutant cancers (Chiti & Dobson, 2017; Iwahashi et al., 2020). Other recent studies have demonstrated the accumulation of misfolded proteins in PE placentas, suggesting that PE also belongs to a class of protein conformational diseases (Cater et al., 2019; Cheng et al., 2022). Several studies have reported placental deposition of aggregates of Aβ and transthyretin, whose mutants cause the most common form of hereditary amyloidosis (Buxbaum, Koziol, & Connors, 2008; Raz, Shiratori, & Goodman, 1970), in PE placentas (Cater et al., 2019; Cheng et al., 2022). However, these studies did not address how these aggregates contribute to poor placentation. Down syndrome is caused by trisomy of chromosome 21, on which APP is located, and is characterized by marked accumulation of Aβ fibrils in the brain (Burger & Vogel, 1973; Oyama et al., 1994; Wisniewski, Wisniewski, & Wen, 1985). The study by Wong et al. reported that overexpression of APP in BeWo cells inhibited syncytialization (Wong, Cheung, Ip, Ngan, & Cheung, 2018), but the effect of Aβ fibrils on syncytialization of BeWo cells was not investigated. Our study provides the evidence and mechanism that Aβ fibrils indeed inhibit STB formation and may contribute to the PE pathology.

Because placental hypoxia plays an important role in the pathophysiology of PE (Tong et al., 2022), we hypothesized that hypoxia in the PE placenta increases Aβ production and that Aβ deposited in the placenta affects syncytialization Here, pre-treatment of BeWo cells and human CTBs with Aβ42 fibrils significantly reduced β-hCG secretion and induction in the medium and syncytin-1 expression, indicating that Aβ42 fibrils inhibited syncytialization and STB formation by CTBs. Knockdown of ZO-1 decreased cell-cell fusion and subsequent trophoblastic differentiation (Pidoux et al., 2010), and reduced cell surface expression of E-cadherin led to dysfunctional cell-cell adhesion and disturbed syncytialization (Iwahashi et al., 2019). The proper function of cell-cell adhesion proteins depends on their appropriate membrane localization, but is not reflected in lower protein levels (Tsukita, Furuse, & Itoh, 2001; Yap & Kovacs, 2003). Aβ42 assemblies disrupted the membrane localization of tight junctional proteins, including ZO-1, by inducing autophagy in murine cerebral capillary endothelial cells (Kook et al., 2012), and E-cadherin turnover was regulated via the endocytosis and autophagic pathway (Santarosa & Maestro, 2021). Therefore, BeWo cells treated with Aβ42 fibrils showed a significant increase in LC3 levels, suggesting that increased autophagy led to reduced ZO-1 and E-cadherin. Pre-treatment of CTBs with Aβ42 fibrils induced loss of membrane localization of cell-cell adhesion proteins, which are essential for CTB syncytialization. These results suggest that Aβ deposition is increased in the hypoxic PE placenta and that Aβ fibrils enhance autophagy of CTB, thereby disrupting the proper membrane localization of cell adhesion-related proteins such as ZO-1 and E-cadherin and inhibiting syncytialization, which may be a cause of placental dysfunction and PE.

Our results suggest that Aβ productions by CTBs are at least partially regulated by hypoxia and HIF1-α, however, other factors that were implicated in increased Aβ production, such as endoplasmic reticulum stress (Fu, Zhao, Wang, & Zhu, 2015; Jung et al., 2020; Lian et al., 2011) and oxidative stress (Li et al., 2004; Raijmakers, Dechend, & Poston, 2004), may be involved in Aβ production by CTBs in the late stage of gestation. The stage at which Aβ begins to deposit in PE placentas is still unclear. Defective spinal artery remodeling can result in chronic and pathogenic hypoxia, which would lead to an increase in Aβ42 production and Aβ deposition. Additional studies are needed to elucidate the significance of Aβ metabolism and deposition in the pathology of the human placenta. Cheng et al. reported that hypoxia and subsequent induction of endoplasmic reticulum stress induced abnormal accumulation of TTR aggregates in the placental junctional zone (Cheng et al., 2022), however, the contribution of TTR aggregates to poor placentation was not fully clarified. Although our results clearly indicate the detrimental role of Aβ fibrils in the maintaining pregnancy, future studies are also needed to elucidate the pathological roles of different protein aggregates.

In summary, we showed here that Aβ fibrils are detrimental to CTB syncytialization in PE placentas. The current view in AD is that Aβ oligomers are the primary culprit in synaptic defects (Vassallo, Ongeri, & De Simone, 2023). The study of Aβ oligomers in human placenta will be an important future research topic.

## MATERIALS AND METHODS

### Materials

Aβ peptides (human, 1-42) were purchased from Peptide Institute (Osaka, Japan). Rabbit polyclonal anti-β-actin and a rabbit polyclonal anti-microtubule-associated protein 1A/1B-light chain 3 (LC3) antibodies were purchased from Medical and Biological Laboratories (Tokyo, Japan). Rabbit monoclonal anti-HIF1α antibody was purchased from Cell Signaling Technology (Danvers, MA, USA), and a mouse monoclonal anti-BACE1 antibody was from R&D Systems (Minneapolis, MN, USA). Rabbit polyclonal anti-β-hCG antibody was obtained from Proteintech (Rosemont, IL, USA). Rabbit polyclonal anti-ERVWE1/HERV/Syncytin antibody, a rabbit polyclonal anti-ZO-1 tight junction protein antibody, and a mouse monoclonal anti-E-cadherin antibody were purchased from Lifespan Biosciences (Seattle, WA, USA), Abcam (Cambridge, UK), and BD Biosciences (Franklin Lakes, NJ, USA), respectively. LY2886721, the BACE1 inhibitor, was purchased from Abcam. Roxadustat was purchased from Selleck Chemicals (Houston, TX, USA). Anti-human BACE1 (C) rabbit IgG affinity-purified antibody and anti-human Aβ1-42 rabbit IgG affinity-purified antibody were purchased from Immuno-Biological Laboratories (Gunma, Japan). Anti-amyloid β40 mouse monoclonal antibody BA27 was obtained from Wako Pure Chemicals (Osaka, Japan), and the 22C11 anti-APP antibody was from Thermo Fisher Scientific (Waltham, MA, USA). β-Secretase-cleaved Nt-specific rabbit polyclonal anti-Aβ antibody (β001) was established as previously described (Lippa et al., 1999). RB4CD12 anti-S-domain antibody was kindly provided by Dr. Toin H. van Kuppevelt (Nijmegen Center for Molecular Life Sciences, Radboud University Nijmegen Medical Center, Nijmegen, The Netherlands) (Dennissen et al., 2002).

### Human tissue collection

This study was approved by the Ethics Committee of Wakayama Medical University (authorization no. 1690) and was conducted according to the tenets of the Declaration of Helsinki. All patients gave written informed consent for the use of tissue samples in this study. Third-trimester human placentas were collected after vaginal deliveries or cesarean sections. HDP (hypertensive disorders of pregnancy) is defined as hypertension (blood pressure ≥ 140/90 mmHg) during the pregnancy. PE is one type of HDP and is accompanied by one or more of the following new-onset conditions at or after 20 weeks of gestation: proteinuria (≥300 mg protein/24 hours); other maternal organ dysfunctions including liver involvement without any underlying diseases; progressive kidney damage; stroke; neurological complications such as clonus, eclampsia, visual field disturbance, and severe headache except for primary headache; hematological complications such as thrombocytopenia caused by an HDP-related platelet count lower than 150,000/μl, disseminated intravascular coagulation, and hemolysis; and uteroplacental dysfunction such as fetal growth restriction, abnormal umbilical artery Doppler waveform results, and stillbirth. All the above-mentioned symptoms and signs become normal by 12 weeks’ postpartum. The eligibility of the PE cases was determined according to the diagnostic criteria of the International Society for the Study of Hypertension in Pregnancy. Cases involving multiple pregnancies, fetal chromosomal abnormalities, and fetal anomalies were excluded. For immunohistochemical analysis, placental tissue samples from PE patients and gestational age-matched controls were washed in ice-cold phosphate-buffered saline (PBS) (pH 7.2) before fixation with 4% paraformaldehyde in PBS (*n* = 5 for normal pregnancy and *n* = 5 for PE).

### Immunohistochemistry

Paraffin-embedded placenta blocks were cut into 3-μm-thick sections, after which they were deparaffinized and rehydrated. Epitopes were then retrieved via heat by boiling sections in a pressure cooker in citrate-based Antigen Unmasking Solution (pH 6.0; Vector Laboratories, Newark, CA, USA) for 20 min. Sections were then incubated with the β001 anti-Aβ antibody (1:1000) and RB4CD12 (1:100), or an anti-HIF-1α antibody (1:100) and an anti-BACE1 antibody (1:50), followed by use of an Alexa Fluor 488-conjugated polyclonal goat anti-mouse IgG (1:300; Thermo Fisher Scientific), a Cy3-conjugated polyclonal goat anti-rabbit IgG, an Alexa Fluor 488-conjugated polyclonal goat anti-mouse IgG (1:400; Thermo Fisher Scientific), or a Cy3-conjugated monoclonal anti-vesicular stomatitis virus G glycoprotein (1: 300; Sigma, St. Louis, MO, USA). Sections were mounted with Vectashield mounting medium with 4’,6-diamidino-2-phenylindole (DAPI) (Vector Laboratories) and were examined with an LSM700 confocal microscope (Carl Zeiss, Jena, Germany). For quantification, 5 regions of interest (ROIs; 16 µm × 16 µm) were set on villi in each placenta tissue, and mean intensities were determined by using ZEN 3 blue edition (Carl Zeiss). β001- and RB4CD12-positive area were determined by using the Coloc. Tools of ZEN 3.

### Cell culture

Human choriocarcinoma-derived BeWo cells that are widely used as a model of trophoblast syncytialization were purchased from the Japanese Collection of Research Bioresources (Osaka, Japan). BeWo cells were maintained in minimal essential medium-α (MEM-α) (Wako Pure Chemicals) supplemented with 15% FBS and the antibiotic mixture. Cells were cultured at 37°C in an atmosphere of 5% CO_2_ and 95% air.

### Hypoxic treatments

Hypoxic conditions (2% O_2_) were established by using the BIONIX-1 hypoxic culture kit (Sugiyamagen, Tokyo, Japan) containing an AnaeroPack-Anaero 5% system (an oxygen absorber; Mitsubishi Gas Chemical, Tokyo, Japan), an OXY-1 oxygen monitor (JIKCO, Tokyo, Japan), an AnaeroPouch (Mitsubishi Gas Chemical), and plastic clips for sealing pouches. Cells were grown in 6-cm dishes or 6-well, 12-well, or 24-well plates, and an OXY-1 oxygen monitor and an oxygen absorber were arranged in a pouch and the left open side was sealed with a clip. When the O_2_ concentration in the pouch reached 2%, the pouch between the culture dish or plate and the oxygen absorber was sealed with another clip to prevent additional oxygen absorption, and the pouch was maintained at 37°C.

### Aβ fibril preparation

Aβ fibrils were prepared as previously described (Z. Zhang et al., 2017). Briefly, chemically synthesized Aβ peptide (human, 1–42; Peptide Institute) was dissolved in 0.1% NH_3_ to prepare a 1 mM stock solution. Stock solution was diluted in PBS to 200 μM and was incubated at 37°C for 5 days to prepare Aβ fibrils. The formation of Aβ fibrils was confirmed by measuring the fluorescence intensity of thioflavin T and by transmission electron microscopy analysis (Supplementary Fig. S4).

### Western blot analysis

Aβ generated by BeWo cells was analyzed by means of Western blotting. BeWo cells were cultured in Opti-MEM medium with 2% FBS with or without the BACE1 inhibitor (LY2886721) for 24 hours under normoxic conditions (20% O_2_) or hypoxic conditions (2% O_2_), respectively. CM samples were then collected and centrifuged at 2000*g* for 30 min to remove debris, after which 100% trichloroacetic acid (TCA) in PBS was added to obtain a final concentration of 10%, and the mixture was incubated for 30 min at 4°C, followed by centrifugation at 12,000*g* for 10 min at 4°C. Precipitates were dissolved in UTB (9 M urea, 2% TritonX-100, 5% 2-mercaptoethanol) and were sonicated on ice. The 2× sodium dodecyl sulfate-polyacrylamide gel electrophoresis (SDS-PAGE) sample buffer (4% sodium dodecyl sulfate, 12% 2-mercaptoethanol, 0.1 M Tris, 20% glycerol, and 0.01% bromophenol blue) was then added to the samples, which were sonicated on ice to prepare PAGE samples that were subjected to Western blotting as described below.

The effects of Aβ fibrils on CTB differentiation were investigated by analyzing β-hCG production and syncytin-1 expression via Western blotting. Fsk was used as an inducer of syncytialization of BeWo cells. BeWo cells were pre-treated with Aβ42 fibrils (10 μM) in serum-free Opti-MEM for 12 hours and then stimulated with Fsk (50 μM) for 48 hours. To analyze ZO-1, E-cadherin, and LC3 expression, BeWo cells were cultured in serum-free Opti-MEM in the presence or absence of Aβ42 fibrils (10 μM) for 24 hours. To investigate β-hCG production and secretion, CM samples were collected and centrifuged at 2000*g* for 30 min to remove debris. CM obtained was mixed with 2× SDS-PAGE sample buffer and heated at 95°C for 5 min. To determine protein levels of β-hCG, syncytin-1, ZO-1, E-cadherin, and LC3, cells were fixed with 10% TCA and whole cell lysates were prepared as described above. Samples were then subjected to SDS-PAGE with 5% to 20% gels (Wako Pure Chemicals) and were transferred to polyvinylidene difluoride membranes (Millipore, Burlington, MA, USA). To analyze Aβ production, PAGE samples were subjected to NuPAGE with NuPAGE MES SDS Running Buffer and NuPAGE 4–12%, Bis-Tris gels (Thermo Fisher Scientific). Membranes were blocked in the EzBlock Chemi blocking solution (ATTO, Tokyo, Japan) or 5% bovine serum albumin (Proliant Biologicals, Ankeny, IA, USA) and 0.1% Tween 20 in Tris-buffered saline at room temperature for 1 hour, after which they were incubated with the anti-hCG β antibody (1:1000), the anti-syncytin-1 antibody (1:1000), the anti-ZO-1 antibody (1:1000), the anti-E-cadherin antibody (1:1000), or the anti-LC3 antibody (1:1000), followed by a preabsorbed horseradish peroxidase-conjugated anti-rabbit or anti-mouse IgG (1:10,000, Jackson ImmunoResearch Laboratories, West Grove, PA, USA). Signals were visualized by using ImmunoStar LD Chemiluminescence Reagent (Wako Pure Chemicals) and an Amersham ImageQuant 800 system (Cytiva, Marlborough, MA, USA). To measure Aβ, membranes were incubated with the β001 rabbit polyclonal anti-Aβ (Nt) antibody (1:10,000) followed by a preabsorbed horseradish peroxidase-conjugated anti-rabbit (1:10,000; Jackson ImmunoResearch Laboratories).

### Immunocytochemistry

BeWo cells were cultured on cover glasses and treated with Aβ fibrils (10 μM) for 24 hours at 37°C, after which they were fixed in 4% paraformaldehyde in PBS at room temperature for 20 min. After the cells were washed three times with PBS, they were blocked and permeabilized with 20% Animal-Free Blocker (Vector Laboratories) containing 0.05% saponin (Wako Pure Chemicals) in PBS for 20 min at room temperature. The samples were then incubated with the anti-ZO-1 antibody (1:100) or the anti-E-cadherin antibody (1:200) followed by incubation with Alexa Fluor 488-conjugated polyclonal goat anti-rabbit IgG (1:300) or Alexa Fluor 568-conjugated polyclonal goat anti-mouse IgG (1:300). Specimens were mounted with the Vectashield mounting medium with DAPI and were examined with an LSM700 confocal microscope.

### Isolation of human trophoblasts

All patients provided written informed consent for the use of tissue samples. Villous CTBs were isolated as previously described (Simon et al., 2017). Placental tissues from 4 normal full-term pregnancies were obtained by elective cesarean section before the onset of labor. Villous tissues were washed three times with PBS, and chorionic and basal plates and large vessels were removed, after which tissues were again washed with PBS and minced. We digested villous tissues with trypsin (0.25%, Thermo Fisher Scientific) and DNase (0.03 mg/ml, Merck), and the supernatants obtained were filtered by using 100-µm cell strainers (SPL Life Sciences, Gyeonggi-do, Korea). FBS samples (5 ml) were added slowly to the bottoms of the tubes containing the supernatants, which were centrifuged at 1250*g* for 15 min at room temperature. After we gently removed the supernatants, we resuspended cell pellets that contained red blood cells and CTBs in RPMI 1640 medium supplemented with 10% FBS. Suspensions were layered on top of a 20–60% Percoll (Cytiva, Marlborough, MA, USA) gradient in Hanks’ Balanced Salt Solution (Thermo Fisher Scientific). After centrifugation at 1250*g* for 20 min at room temperature, the CTB layers were collected. RPMI 1640 medium supplemented with 10% FBS was added to the CTB layers and subjected to additional centrifugation at 1250*g* for 15 min at room temperature. Isolated primary human CTBs were seeded at a density of 1.0 × 10^6^/4 wells in a 12-well plate and grown in Trophoblast Medium (ScienCell Research Laboratories, Carlsbad, CA, USA) supplemented with 5% FBS and an antibiotic mixture containing penicillin and streptomycin (Life Technologies).

Expression of APP, expression of BACE1, and Aβ production in human primary-cultured CTBs were confirmed by using Western blotting with the 22C11 anti-APP antibody, the BACE1 antibody, and the β001 antibody, respectively. The effects of Aβ fibrils on CTB differentiation were analyzed by means of Western blotting with the anti-β-hCG antibody and the anti-syncytin-1 antibody as described above except that primary CTBs were pre-treated with Aβ fibrils (10 μM in Opti-MEM) for 24 hours and then incubated in fresh Opti-MEM with 10 μM Aβ fibrils for an additional 48 hours. The effects of Aβ fibrils on the expression and subcellular localization of ZO-1 and E-cadherin were immunocytochemically analyzed as described above, except that primary CTBs were cultured on poly-L-lysine-coated cover glasses for 9 hours and treated with Aβ fibrils (10 μM) for 18 hours.

### Analysis of the effects of HIF1-α on BACE1 expression

For this analysis, we used roxadustat (Selleck Chemicals), which inhibits prolyl hydroxylase and stabilizes HIF as an HIF1-α stabilizer. BeWo cells were treated with roxadustat (0, 5, 10, or 25 μM) in MEM-α medium containing 15% FBS for 16 6 hours, after which whole cell lysates were prepared and subjected to Western blotting with the anti-HIF-1α antibody (1:1000) and the anti-human BACE1 (C) antibody (1:50), respectively, as described above.

### Statistical analysis

For statistical analysis, we used an ordinary one-way analysis of variance with the post hoc Dunnett, Bonferroni, or Tukey test, or unpaired Student’s *t* test with Prism software (GraphPad Software, Version 7.04, San Diego, CA, USA). Mann Whitney test was used for statistical analysis of the quantification in immunohistochemistry. Results were said to be significant when *P*-values were less than 0.05.

## Author contributions

K. Nishioka, M.I., N.I., M.A., M. Fukushima., N.M., M. Fujino., and K. Nishitsuji performed the experiments. K. Nishitsuji and N.I. conceived the study and designed the experiments. K. Nishioka and K. Nishitsuji interpreted the data and wrote the paper. M.M., Y.H., and K.I. analyzed the human tissues. K.U. contributed experimental materials and tools and reviewed the manuscript. T.T. and Y.I. contributed experimental materials and tools. K. Nishitsuji takes full responsibility for the entire manuscript. All authors reviewed the results and approved the final version of the manuscript.

## Acknowledgement

Thanks are due to Dr. Hiroshi Mori at Nagaoka Sutoku University for his continuous encouragement.

## Data Availability Statement

All data are included in the manuscript and/or supporting information.

