## Supplementary Materials for "Amyloid-β fibrils accumulated in preeclamptic placentas suppress syncytialization of cytotrophoblasts"

**Supplementary Methods**

**Thioflavin T (ThT) Fluorescence Assay and and Transmission electron microscopy (TEM)**

ThT (10 μM) in an Aβ42 fibril solution (200 μM in PBS, pH 7.4) was exited at 445 nm and the fluorescent intensity was recorded from 470 nm to 600 nm by using a MTP-900Lab microplate reader (Hitachi High-Tech Science, Tokyo, Japan). For TEM analysis, Aβ42 fibrils (200 μM in PBS) were spread on carbon film-coated copper grids and negatively stained twice with 2% uranyl acetate for 1 min. The grids were then examined under a JEM-1400Plus transmission electron microscope (JEOL, Akishima, Japan) with an acceleration voltage of 100 kV. Digital images (3296 × 2472 pixels) were obtained by using a charge-coupled device camera (EM-14830 RUBY2, JEOL).

**Analysis of E-cadherin mRNA**

To analyze the expression of E-cadherin, BeWo cells were cultured in serum-free Opti-MEM in the presence or absence of Aβ42 fibrils (10 μM) for 24 hours. Total RNA was obtained by using the TRIzol reagent (Thermo Fisher Scientific). RT-qPCR was carried out with the CFX96 Touch Real-Time system (Bio-Rad Laboratories, Hercules, CA) and the iTaq universal Universal SYBR Green one-step kit (Bio-Rad). RT-qPCR experiments were performed in triplicate. Data were processed by using the Bio-Rad CFX Manager version 3.1 (Bio-Rad Laboratories), and expression levels were calculated via the comparative ΔΔCt method by using GAPDH as the reference gene.

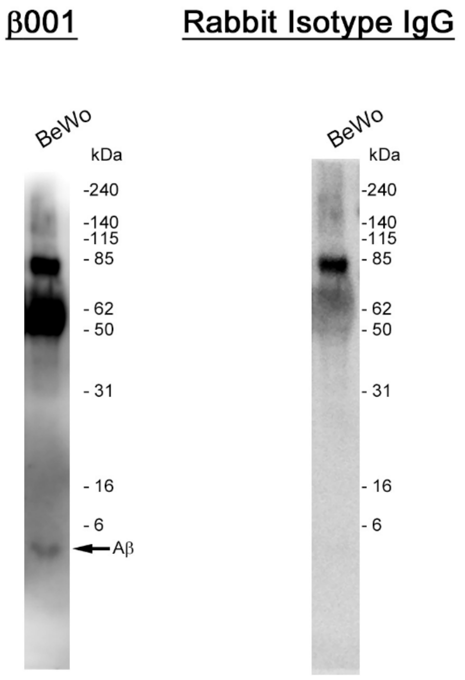


**Figure S1.** Conditioned media from BeWo cells were analyzed by β001 and the membrane was re-probed with an isotype rabbit IgG. High molecular weight bands were observed in both immunoblots, indicating that these bands are non-specific.

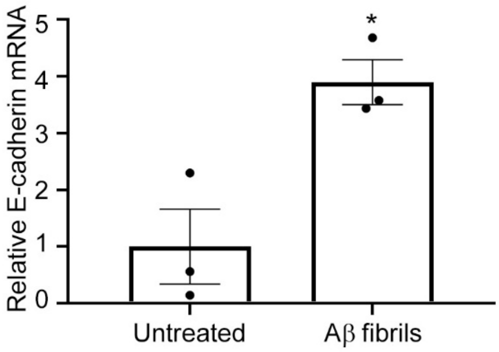


**Figure S2.** Aβ fibril treatment enhanced E-cadherin mRNA expression in BeWo cells. BeWo cells were treated with Aβ1-42 fibrils (10 μM) in serum-free Opti-MEM for 24 hours. E-cadherin transcriptional levels were analyzed by means of semi-quantitative RT-PCR. Data are means ± SEM ($n$ = 3). GAPDH served as the reference gene. The sequences of the primers used were as follows: E-cadherin, forward: CAAATCCAACAAAGACAAAGAAGGCAA, reverse: ATGACAGACCCCTTAAAGACCTCCT; GAPDH, forward: GAGTCAACGGATTTGGTCGT, reverse: GACAAGCTTCCCGTTCTCAG. *$P$ < 0.05.

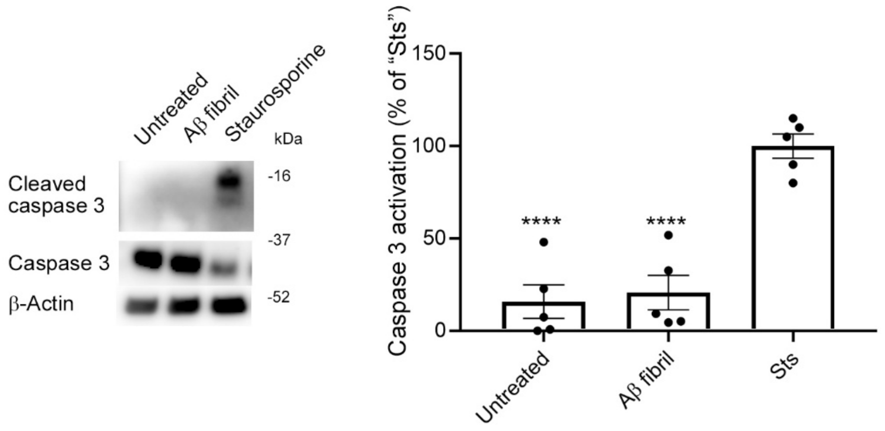


**Figure S3.** No activation of caspase 3 by Aβ fibrils in BeWo cells. BeWo cells were treated with Aβ fibrils (10 μM, 24 h) or staurosporine (Sts, Selleck Chemicals, 1.0 μM, 3 h), after which caspase activation was analyzed by means of Western blotting with a rabbit monoclonal anti-cleaved caspase 3 antibody and a rabbit polyclonal anti-caspase 3 antibody (Cell Signaling Technology). β-Actin is used as a loading control. Data are means ± SEM ($n$ = 5). ****, *$P$ < 0.0001 versus "Sts".

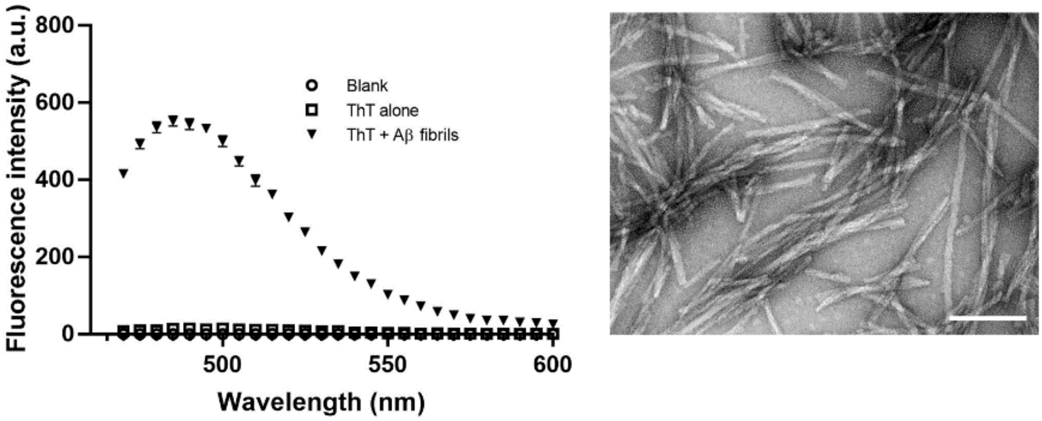


**Figure S4.** ThT assay (left) and TEM analysis (right) of Aβ fibrils (200 μM). "Blank": PBS in the absence of ThT and Aβ fibrils; "ThT alone": ThT (10 μM) in the absence of Aβ fibrils. Scale bar in the TEM image: 100 nm.
